# Description and lessons learned from the 2014 Dengue outbreak in Dar es Salaam, Tanzania. Knowledge, attitudes and bite prevention practices among those with confirmed Dengue

**DOI:** 10.1101/567396

**Authors:** Daniel Msellemu, Tegemeo Gavana, Hassan Ngonyani, Yeromin P. Mlacha, Prosper Chaki, Sarah J. Moore

## Abstract

**Background:** The frequency and magnitude of Dengue epidemics have increased dramatically in the past 40 years throughout the tropics largely due to unplanned urbanization, globalization and lack of effective mosquito control. Dar es Salaam, Tanzania has recently experienced Dengue outbreaks that occur with increasing frequency. Currently, only one serotype is recorded. Without adequate vector monitoring and control, it is certain that further outbreaks will occur.

**Methods/Findings:** A retrospective study followed 100 individuals with confirmed Dengue fever in Kinondoni, Dar es Salaam during the 2014 outbreak. Houses were inspected for mosquito breeding sites and gathered information on Socio-economic Status (SES) and Dengue prevention knowledge.

Higher SES tertile had the most Dengue cases: 53 (55%) followed by medium and lower SES with 33 (34%) and 11 (11%) respectively. The highest number of mosquito breeding sites was also found in higher SES households. Kinondoni wards of Manzese, Mwananyamala, Tandale and Mabibo had the highest number of confirmed cases: 18, 13, 13 and 9 respectively. Each ward has large marketplaces, which may have aided dissemination of transmission to other areas.

The population remains poorly informed about Dengue transmission: 22% of respondents said Dengue is spread from person to another, 30% did not think mosquitoes spread Dengue and 60% heard about Dengue while in hospital. Knowledge of bite prevention was poor; Dengue mosquito bites outside of sleeping hours but 84% of Dengue patients said that using bednets would prevent vector bites.

**Conclusion:** Affluent households are likely to be reservoirs of Dengue vectors having more breeding sites and Dengue cases. Mobile phones whose ownership is high across all social classes seem to be a better tool to communicate information about Dengue. The study established a habitat suitability score, a tool to be used for learning and estimate breeding habitat capacity to be used for vector control before rains begin.

**Author’s Summary:** Dengue fever is a viral infection transmitted by Aedes (Stegomyia) mosquitoes causing a flu-like illness that may develop into severe complications such as Dengue haemorrhagic fever and Dengue shock syndrome if the patient contracts two viral serotypes concurrently. There is currently no antiviral treatment or vaccines available against Dengue. Environmental vector control and mosquito bite prevention remain essential to prevent transmission. Due to globalisation and rapid urban expansion, Dar es Salaam is experiencing regular Dengue outbreaks. Without adequate vector control and public awareness, it is certain that these will continue to re-occur.

The study presents factors associated with the outbreak in 2014. Rich households have more places for mosquitoes to breed with 54% found in these households and the majority of Dengue cases 55% came from higher SES groups that represented a greater proportion of cases than lower and middle socioeconomic groups, combined. The public was ill-informed about Dengue fever: 84% think bed nets can prevent Dengue, and 60% of the patients only became aware of Dengue while in the hospital with the illness. The study established a habitat suitability score, a tool to be used to estimate breeding habitat capacity before rains begin. Scattered containers especially tyres remain ideal breeding sites. The study highlights the need for waste management to avert future outbreaks.

## Introduction

In 2014, a Type 2 Dengue fever outbreak in Tanzania spread to seven regions on the mainland and two regions in Zanzibar [1]. On mainland Tanzania, there were 1,017 confirmed cases from a total of 2,121 suspected cases including 4 deaths. Zanzibar had 1 confirmed case out of 8 suspected cases and no deaths. Ninety-nine per cent (99%) of the cases of the mainland were reported from the following three districts of Dar es Salaam: Kinondoni, Temeke, and Ilala. Of the four fatal cases, 1 had presented with Dengue Haemorrhagic Fever (DHF) and 1 with multiple organ failure [2].

Dengue is currently the most widespread arboviral (insect-transmitted virus) disease of humans. The frequency and magnitude of Dengue epidemics have increased dramatically in the past 40 years as the four virus serotypes and the mosquito vectors have both expanded geographically in the tropics and subtropics. The principal factors that have contributed to this emergence of epidemic Dengue are 1) unplanned urbanization, 2) globalization and 3) lack of effective mosquito control [3]. Tanzania’s urban population is rapidly expanding with 30% of the population now living in Urban areas [4], with 4.3 million residing in Dar es Salaam which also hosts Tanzania’s largest airport and handles 90% of shipping cargo. In addition, Tanzania has strong trade and economic links with many Dengue endemic countries in South-east Asia providing a route for the introduction of new serotypes.

In Dar es Salaam there is widespread unplanned urbanisation without waste management. This results in extensive areas of household waste such as plastic containers that provide ample breeding sites for vector mosquitoes [5].

The reported incidence of Dengue has increased worldwide in recent decades, but little is known about its incidence in Africa. During 1960–2010, a total of 22 countries in Africa reported sporadic cases or outbreaks of Dengue [6]. However, Dengue is being more frequently reported. The presence of disease and high prevalence of antibodies to Dengue virus in available serologic surveys suggest endemic Dengue virus infection in many parts of Africa, including Tanzania [7]. Dengue is likely under-recognized and underreported in Africa because of low awareness by health care providers, other prevalent febrile illnesses, and lack of diagnostic testing and systematic surveillance. While Dar es Salaam has in the recent past experienced Dengue outbreaks, without adequate vector control it is certain that these will re-occur and there is a possibility of introduction of other Dengue serotypes and other arbovirus transmissions in the future with 22 million people living in areas of Tanzania suitable for transmission of these diseases [8].

### Dengue Control

There is currently no antiviral treatment or vaccines against Dengue although vaccines are under development [9]. Thus preventing or control of Dengue virus transmission depends entirely in controlling the mosquito vectors or interruption of human-vector contact [10]. Transmission control activities should target the mosquito vector *Aedes aegypti* in the household and immediate vicinity as well as other settings where human-vector contact occurs, such as schools, hospitals, markets and workplaces. It has been shown that efforts to control *Ae. aegypti* are only sustainable through conscious, systematic, proactive and preventive actions carried out by persons and groups. Cuba has remained Dengue free for 30 years by using a “set of measures directed towards the detection and elimination of possible mosquito breeding places. It hinges on weekly self-directed inspection by families and workers in their homes and workplaces” [11]. Environmental management seeks to change the environment in order to prevent vector breeding by destroying, altering, removing or recycling containers that provide mosquito-breeding habitats and should be the mainstay of Dengue vector control. *Ae. aegypti* uses a wide range of confined larval habitats, both man-made and natural[12]. Some man-made container habitats produce large numbers of adult mosquitoes, whereas others are less productive. Consequently, control efforts should target the habitats that are most productive and hence epidemiologically more important rather than all types of containers, especially when there are major resource constraints [10]. With this in mind, we followed up patients with confirmed cases of Dengue during the 2014 outbreak in order to characterize possible risk factors for Dengue during a Dengue fever outbreak in Dar es Salaam “Fig.1”. We followed up confirmed Dengue cases among Tanzanians by visiting their homes and assessing breeding sites and testing a tool to assess Dengue vector breeding habitat potential during the dry season (habitat suitability-score) that could be used for training larval control staff to prevent epidemics. In addition, we conducted a short questionnaire on Dengue patients’ daily activities and travel behaviour around the time of infection to characterise behaviour that might increase the risk of exposure in comparison to the average for the location. The epidemiological and entomological characterisation of the Dengue outbreak is of high importance to the Tanzanian Government to identify strategies needed for the prevention of future outbreaks.

**Fig 1.**
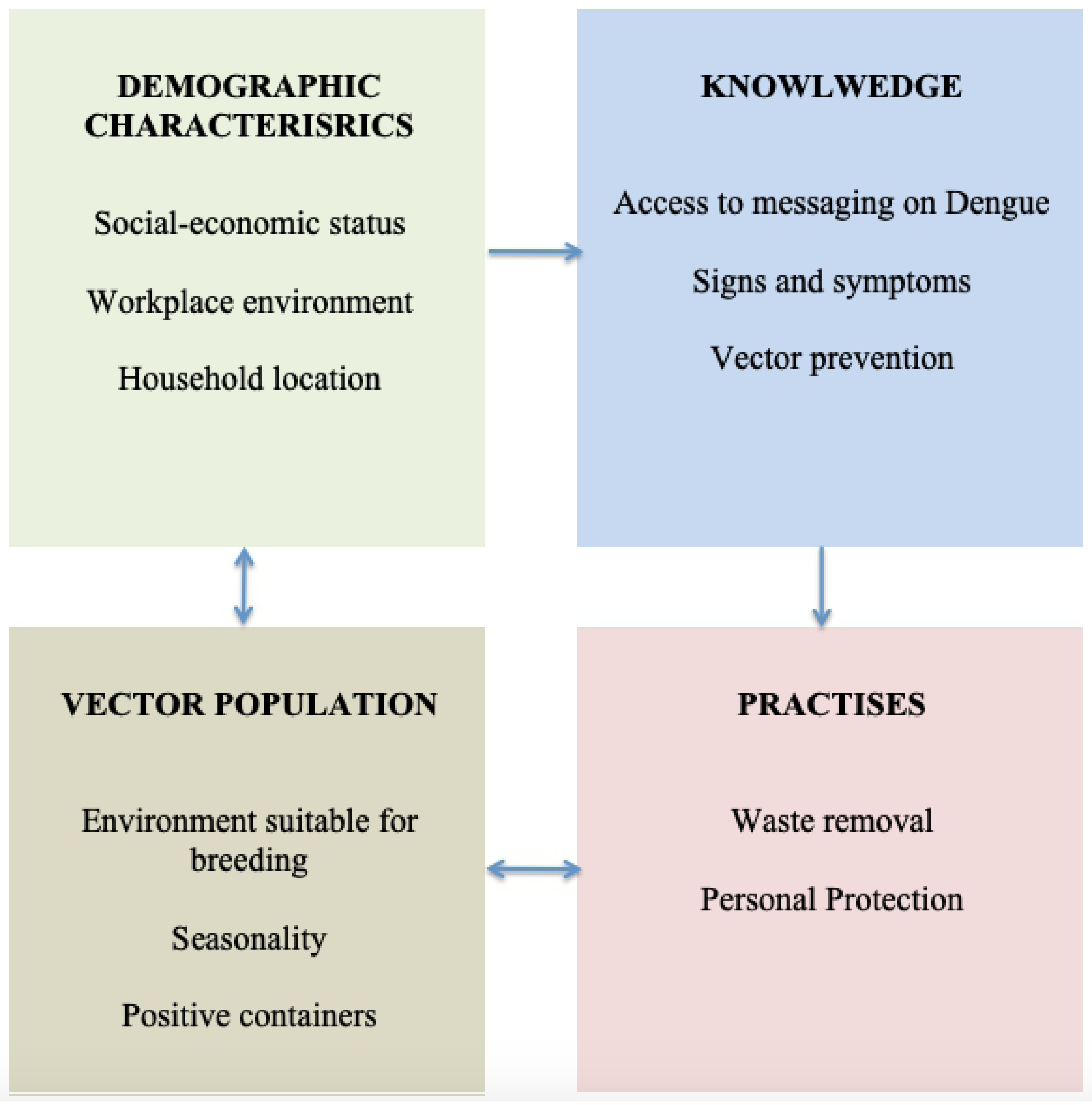
Factors involved in the control or spread of Dengue through Human and *Aedes aegypti* interaction.

## Methodology

### Study Area and Population

A retrospective study following up individuals with confirmed Dengue cases from Mwananyamala hospital, which occurred between January and July 2014, was performed in Kinondoni district from 1st to 28^th^ July 2014. Kinondoni is at the northeastern part of Dar es Salaam city “Fig. 2” with an area of 531 km2, 2,497,940 population and a population density of 1,179 persons per square kilometre [13]. It has an average annual temperature of 25°C and precipitation of 132.9 mm in the rainy season (May– October) [13]. The district has a non-continuous water supply (every 2 days), which leads to widespread water storage, and irregular refuse collection that creates man-made container breeding habitats for mosquitoes.

**Fig 2.**
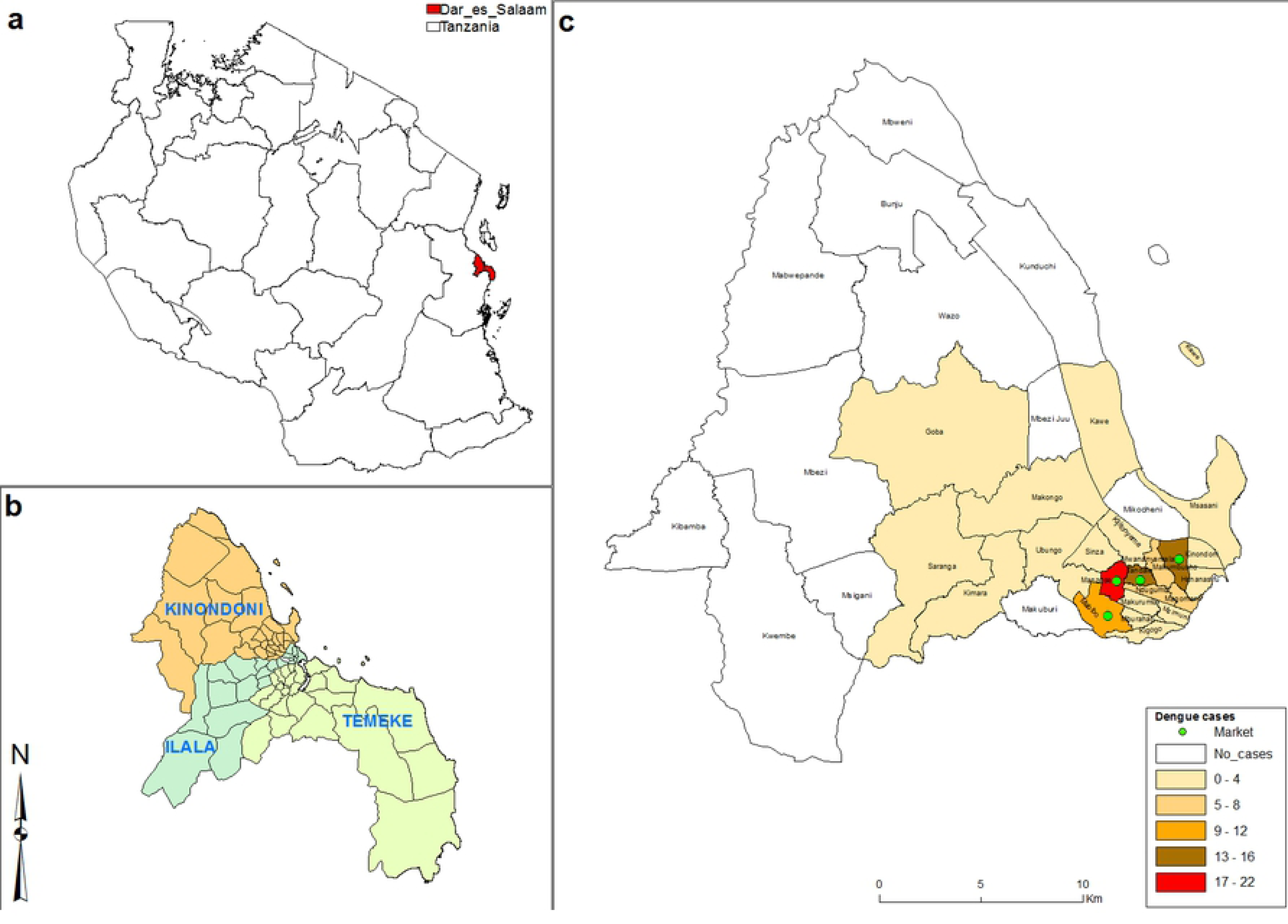
The Map of (a) Tanzania, (b) Dar es Salaam, (c) Kinondoni wards with Dengue cases in 2014.

### Study Site

The “Etiologies of acute febrile illness among adults attending an outpatient department in Dar es Salaam” ClinicalTrials.gov Identifier: NCT01947075 generated a list of 202 Tanzanians with confirmed Dengue Fever from the out-patients department of Mwananyamala district hospital and in other health facilities in Kinondoni District (Sinza hospital, Magomeni healthcare centre, and Tandale dispensary) in Dar es Salaam. From this list, a selection of 100 participants was obtained by computer-generated randomization without repetition. All of the patients had been discharged from the hospital. Only adults of over 18 years of age who were residents of Kinondoni municipal and were sick during the outbreak were interviewed. It should be noted that around 200 cases in non-Tanzanian nationals also occurred but these persons were not followed up as part of this study.

### Data Collection

The study collected information on Dengue patient history before and after the Dengue outbreak. The data collection team visited patients’ homes and upon written informed consent they conducted a short questionnaire on behaviours that increase risks of Aedes mosquito bites and examined the environment looking for breeding sites and then collected larval mosquitoes found in and around the home. Information collected was breeding habitat photos, rainfall data, larvae, disease history, outdoor activities, mosquito bite prevention practices, Dengue awareness and Socio-economic status (SES). If a selected Dengue case was missing we replaced with a randomly chosen case from the remaining half of 100 cases. Data collection took place in one week in order to minimize temporal heterogeneity that could occur due to rainfall, and also to collect data as close to the time of dengue diagnosis to reduce recall bias. The study was not conducted during the peak of Dengue infection due to delays in obtaining ethical approval although the tail end of the epidemic was ongoing.

Twelve Kinondoni Municipal Health Workers were recruited to collect data. They received two weeks of training on informed consent, questionnaires, larval sampling and photography. They are familiar with Kinondoni district as they have continuously been involved in Dar es Salaam Urban Malaria Control Program (UMCP) and are highly experienced in community liaison and larval mosquito sampling.

#### Rainfall Data

In order to establish rainfall patterns and Dengue outbreak, rainfall data from two months before the outbreak December 2013 to July 2014 was obtained from Tanzania Meteorological Agency (TMA). The rainfall data was compared to Dengue fever outbreak records at Mwananyamala Hospital in Kinondoni District.

#### Photography and Habitat suitability Score

All possible breeding habitats both indoors and outdoors, wet or dry were inspected and assessed for the presence or absence of mosquito larvae and pupae. Areas that qualified these criteria were photographed as possible breeding habitat for Aedes mosquitos [14-16].

Aedes lay eggs during the day in dark-coloured containers with wide openings, in shaded locations and prefer water rich in organic materials such as decaying leaves and algae [14, 15]. Suitable sites include tree-holes, plant axils, and artificial containers, plant pots and vases, discarded tires rainwater tanks, wells, and metal drums, garbage bins and upturned lids, buckets, cups, glasses, jugs, bowls, dishes, trays, drink cans, food containers and pet bowls [17], while old tyres, water storage containers and vegetation were identified to be the most productive breeding habitat in Dar es Salaam [18].

Sony DSC-W530 and Kodak DX 3900 were standard pocket digital cameras used. Photographers were directed to move close to the object (breeding habitat). Photographs of potential breeding habitats were taken without zooming to minimize distortion following a standard operating procedure. Photos were given enumeration number that links the compound and the breeding habitat. Photographs of larval habitats taken during the survey were given habitat suitability scores based on “Table 1” the suitability of Aedes breeding habitat [14]. The household habitat suitability was then analysed by SES.

**Table 1.** *Aedes aegypti* breeding habitat suitability scores among houses with confirmed Dengue cases.

| Constituent | Presence | Score |
| --- | --- | --- |
| **Larval** | *Aedes aegypti* larvae identified | 5 |
|  | Other mosquito larvae identified | 3 |
| | Number of breeding habitat 1/2/3/4/∞ | 1-10 |
| **Water** | Accumulated clean water | 5 |
|  | Water storage jars | 5 |
|  | Pools of water with vegetation | 5 |
| **Shade** | Fully shaded habitats | 5 |
|  | Partially shaded habitats | 3 |
|  | Plant axils/flower pots | 3 |
| **Environmental** | Scatted containers | 5 |
|  | Tires | 5 |
|  | Ability to accumulate water during rain season | 5 |

#### Larval collection

Aedes larvae were collected from different breeding habitats in and around the Dengue patients’ houses to within 30 metres radius. All types of containers were thoroughly checked. To avoid leaving behind Aedes larvae, all larvae of any species found in breeding habitat were collected. The larvae were then stored at IHI insectary in emergence breeders for identification of adults.

### Data Management

Questionnaire data were double entered into an Epi Info template corresponding to the questionnaire with drop-down lists of legal values and exported into STATA 12. Data was cleaned through checking for data distribution, unusual values and outliers. Thereafter data was uploaded onto the IHI data server for archiving. No data of a personal nature was collected and the households were identified by GPS.

### Data Analysis

Data were analysed following a predefined analysis plan by descriptive analysis: cross-tabulation with one-tailed Pearson’s Chi-Square to describe associations between Dengue infection and age, gender, socioeconomic status (calculated by Principal Component Analysis), travel history, recall of government Dengue messages and use of personal protection.

The spatial distribution of dengue cases was mapped by using ArcGIS, version 10.5, CA, USA. Coordinates were obtained from dengue cases households. Study site administrative shapefile boundaries were freely downloaded from National Bureau of Statistics website and used to create a map layer.

PCA and Social Economic Status. Socioeconomic status was calculated using a structured questionnaire to gather data on standard variables about household affluence based on the Tanzania Health and Demographic Survey [19]. Variables from Dengue patients’ households that reflecting the economic ability of the people in the community which were pre-specified and apply over time to the study area [20] were used to create Socio-Economic Status (SES) classes from Principal Component Analysis (PCA) [21, 22]. The index was based on three SES categories instead of five as is usually performed because of the small sample size. Data was compared to a larger comparator group of residents obtained from a separate survey in the area – the ABCDR project [23].

### Ethical considerations

Volunteers were recruited on the written informed consent form. Ethical approval was granted by Ifakara Health Institute (IHI) (IHI/IRB/No: 27 -2014) and the National Institute of Medical Research (NIMR/HQ/R.8a/Vol. IX/1866).

## Results

### Relationship between dengue cases Rainfall and SES

When Dengue fever cases started to appear in January 2014 “Fig. 3” cases in our sample came from households with medium and higher SES until March. Even when cases of low SES started to appear they did so in relatively smaller numbers while the middle and higher SES continued to increase. When the rains were at the highest point in April (15mm average), there were only two cases from lower SES while the middle SES and higher SES had 17 and 12 cases, respectively. When the rains diminished in late May to June and finally in July, the higher SES recorded 45 cases, several folds higher than among lower SES and middle SES with 10 and 24 cases, respectively. The higher SES group continued to have cases even when the rains had stopped.

**Fig 3.**
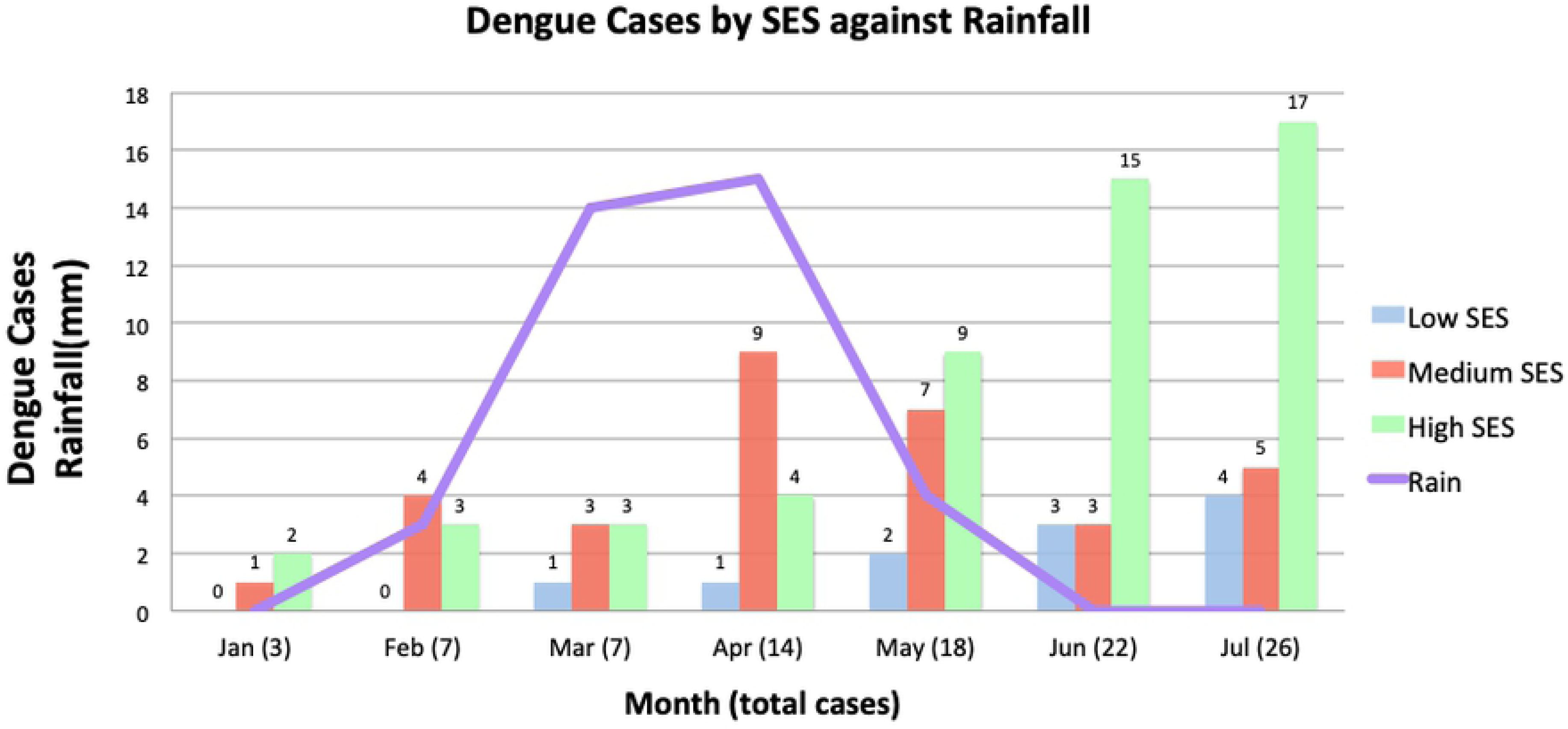
Dengue cases across SES in relationship to rainfall.

There were a total of 202 confirmed Dengue cases in Kinondoni district. A total of 97 (49%) confirmed Dengue patients from the district were interviewed and 97 houses of Dengue patients were visited and evaluated for Aedes breeding places “Table 2”. There were 127 permanent water-holding containers in these houses for domestic use. During the survey that was conducted at the end of the rainy season, 167 potential breeding habitats from 97 houses were identified but only 12 breeding sites that were positive with Aedes. All the *Aedes* were *Ae. aegypti* and no other potential arbovirus vectors such as *Ae. africanus* and *Ae. albopictus* were found.

**Table 2:**
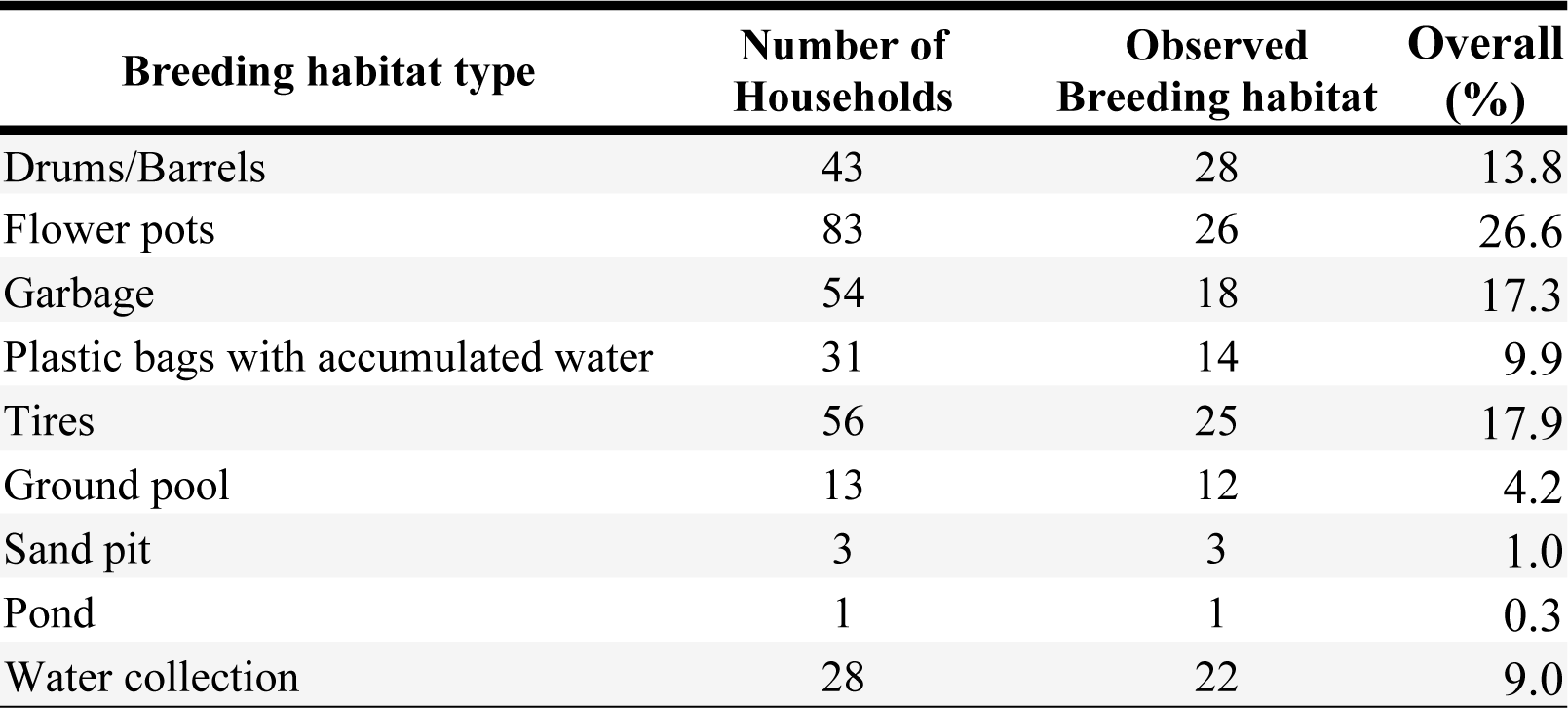
Frequency of common Aedes *aegypti* breeding habitat types across households.

| Breeding habitat type | Number of Households | Observed Breeding habitat | Overall (%) |
| --- | --- | --- | --- |
| Drums/Barrels | 43 | 28 | 13.8 |
| Flower pots | 83 | 26 | 26.6 |
| Garbage | 54 | 18 | 17.3 |
| Plastic bags with accumulated water | 31 | 14 | 9.9 |
| Tires | 56 | 25 | 17.9 |
| Ground pool | 13 | 12 | 4.2 |
| Sand pit | 3 | 3 | 1.0 |
| Pond | 1 | 1 | 0.3 |
| Water collection | 28 | 22 | 9.0 |

The accumulation of Dengue cases in “Fig 4” suggests a propagated outbreak of Dengue virus in the population. This can be seen by a steep increase of Dengue cases and incremental jumps. The jumps reflect the generation of new cases in the population. The two major peaks were in April and July. The accumulation of Dengue cases followed a pattern in which areas close to the market had had a higher number of Dengue cases compared to other parts of Kinondoni district “Fig 2”. Manzese ward had the highest number of cases 18, followed by Mwananymala, Tandale and Mabibo that had 13, 13, and 9 cases, respectively. There was a clear relationship between rainfall data and the Dengue outbreak “Fig3”. The rains started in January and provided breeding sites for Dengue vectors resulting in an increase in Dengue cases and subsequent decrease in cases as the rainy season ended.

### Demographic features of confirmed dengue cases

The average age of people with Dengue was found to be 32 years, which was slightly younger than the average age of the general population (41 years). However, it is important to note that the study was biased because no participants of less than 18 years were followed up for ethical reasons. There was a wide range of ages among Dengue cases with the oldest case being 97 years old. Among this sample from Dar es Salaam the 24 to 34 age-group corresponded to the highest SES and was also the age group with the majority of Dengue patients. More males than females were hospitalised with Dengue 57% (95% CI 46.7-66.7) vs 43% (95% CI 33.3-53.3) “Table 3”.

**Fig 4.**
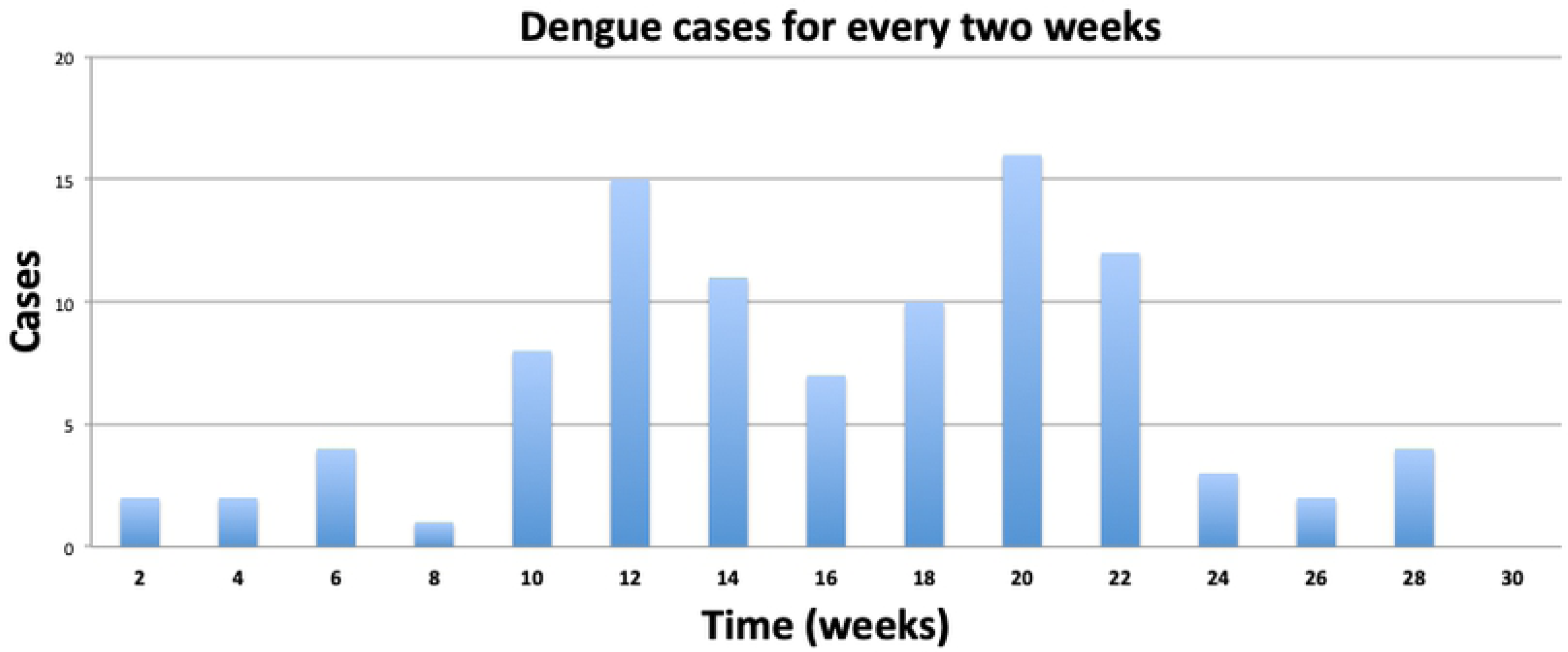
Fortnightly accrual of confirmed Dengue outbreak cases at Mwananyamala hospital clinic in Dar es Salaam Tanzania between January and July of 2014.

**Table 3:** Covariates associated with dengue fever from paired ANOVA. Data are from the Dengue cohort and a concurrent sample from the same area collected by a different study [23] randomised by household to give an estimation of the background SES in the area

| Covariate | Categories | Dengue Patients |  |  | Background SES |  |  | Analysis of variance |  |
| --- | --- | --- | --- | --- | --- | --- | --- | --- | --- |
|  |  | n | (%) | [95% CI] | n | (%) | [95% CI] | Std.Dev | P values |
| Gender | Male | 55 | 57 | [47, 67] | 52 | 30 | [23, 36] | 0.50 | <0.0001 |
|  | Female | 42 | 43 | [33, 53] | 124 | 70 | [64, 77] | 0.43 |  |
| Age categories: Dengue patients and Ordinary population | 18-24 | 31 | 32 | [21, 41] | 20 | 11 | [07, 16] | 0.49 | <0.0001 |
|  | 25-34 | 35 | 36 | [26, 46] | 43 | 24 | [18, 31] | 0.50 |  |
|  | 35-50 | 22 | 23 | [14, 31] | 69 | 39 | [32, 46] | 0.43 |  |
|  | 51 + | 9 | 9 | [03, 15] | 44 | 44 | [19, 31] | 0.37 |  |
| Social Economic Status (SES) comparison between ABCDR and Dengue Patient | Lower SES | 11 | 11 | [05, 18] | 91 | 52 | [44, 59] | 0.31 | <0.0001 |
|  | Medium SES | 33 | 34 | [22, 44] | 47 | 27 | [20, 33] | 0.31 |  |
|  | Higher SES | 53 | 55 | [45, 65] | 38 | 22 | [15, 28] | 0.49 |  |
| Means of health education messaging accessible by the population | Mobile phone | 96 | 99 | [96, 100] | 93 | 53 | [45, 60] | 0.49 | <0.0001 |
|  | Radio | 77 | 79 | [71, 88] | 71 | 40 | [33, 48] | 0.47 | <0.0001 |
|  | Television | 75 | 77 | [69, 86] | 43 | 24 | [18, 31] | 0.47 | <0.0001 |

### Habitat suitability score

Habitat suitability scores “Fig.5” were computed based on breeding habitats “Table 1” around homes with recorded dengue cases. One hundred and twenty-seven pictures were taken from of breeding habitat from 97 houses. The highest habitat suitability score was 24 points and the lowest had 3 points. Mean score was 11.4 points. A household with higher habitat suitability score has a greater potential of harbouring *Aedes aegypti* vectors. It was observed that households with higher SES also have more breeding sites. The average household habitat suitability score increased with SES: Low SES had a mean score of 4 (15.4%), Medium SES: 8 (30.8%) and Higher SES: 14 (53.9%).

**Fig 5.**
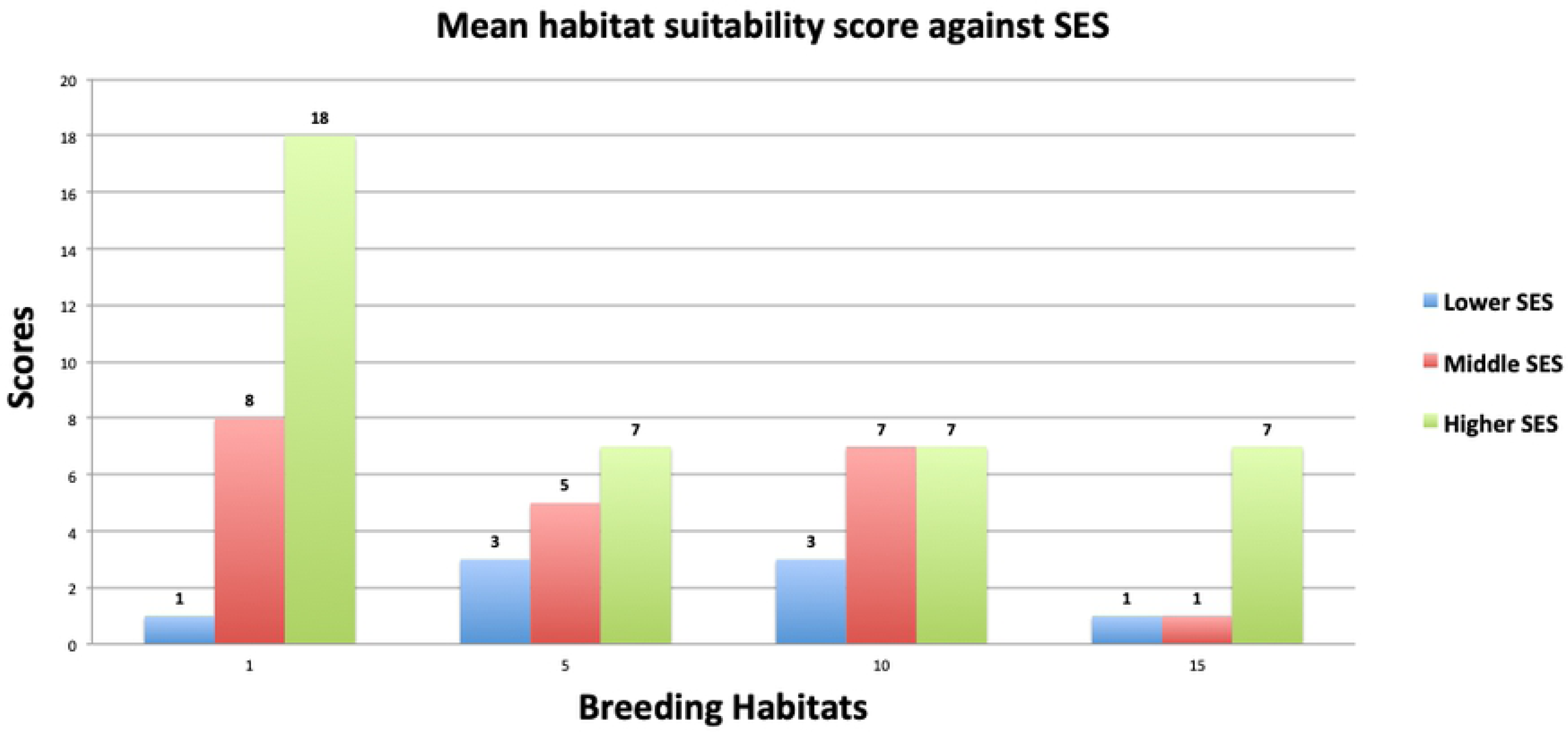
Habitat Suitability Score based on Aedes aegypti bionomics reports of Christophers 1960.

Higher SES households with higher habitat suitability score were associated with water storage and discarded household items such as tyres, buckets, basins, tins and drums. More wealthy houses also had flowerpots and ample space around the home.

### Dengue sign and symptoms

When respondents were asked to identify Dengue signs and symptoms 73 (75%) of them mentioned fever and 82 (85%) mentioned a headache. Other signs and symptoms were: body malaise 62 (63%), joint pain 60 (62%), muscle pain, 26 (27%), eye pain 20 (21%) and vomiting 15 (16%). Other signs and symptoms seldom mentioned were excessive sweating, epistaxis, abdominal pain and diarrhoea.

### Dengue awareness

When asked about where they got information about Dengue fever, most respondents (60%) said at the hospital “Table 4”. The majority of Dengue patients only became aware of Dengue fever when they were already infected. Despite a widespread radio, newspaper and television coverage of the epidemic, radio and television were only mentioned as a source of information about Dengue by 38% and 25% of respondents, respectively, although there was higher radio ownership among dengue patients’ households than the rest of the population. No respondents reported attending a local government meeting about Dengue infection during the outbreak while seeing banners and public announcements were mentioned by 1% of respondents recalling Dengue fever information.

**Table 4.** Knowledge and practice among confirmed Dengue patients from the 2014 Dengue outbreak in Dar es Salaam, Tanzania.

|  | Variable |  | n | % | [95% CI] |
| --- | --- | --- | --- | --- | --- |
| Travel History | Did not Travel |  | 77 | 79 | [71, 86] |
|  | Travelled Tanzania |  | 16 | 16 | [09, 24] |
|  | Travelled Overseas |  | 4 | 4 | [00, 08] |
| Dengue Fever awareness | Radio |  | 37 | 38 | [28, 47] |
|  | Television |  | 25 | 25 | [17, 35] |
|  | Brochure |  | 6 | 6 | [01, 11] |
|  | Public Announcement |  | 1 | 1 | [-01, 03] |
|  | Hospital |  | 59 | 60 | [52, 71] |
| Prevention from mosquito bite | Use of Bed net |  | 84 | 86 | [79, 93] |
|  | Long dress & trousers |  | 16 | 16 | [08, 24] |
|  | Mosquito mesh |  | 26 | 27 | [17, 35] |
|  | Aerosol Sprays |  | 43 | 44 | [34, 54] |
|  | Mosquito coil |  | 6 | 6 | [01, 11] |
|  | Cut grass outdoors |  | 2 | 2 | [-01, 05] |
|  | Fan |  | 20 | 21 | [12, 29] |
|  | Topical repellent |  | 12 | 12 | [06, 19] |
| Dengue Signs and symptoms presented | Fever | Yes | 73 | 75 | [67, 84] |
|  |  | No | 24 | 25 | [16, 34] |
|  | Headache | Yes | 82 | 85 | [77, 92] |
|  |  | No | 15 | 15 | [08, 23] |
|  | Joint pains | Yes | 60 | 62 | [52, 72] |
|  |  | No | 37 | 38 | [28, 48] |
|  | Vomiting | Yes | 15 | 16 | [08, 23] |
|  |  | No | 82 | 84 | [77, 92] |
|  | Eye pain | Yes | 20 | 21 | [12, 29] |
|  |  | No | 77 | 79 | [71, 86] |
|  | Muscle pain | Yes | 26 | 27 | [18, 36] |
|  |  | No | 71 | 73 | [64, 82] |
|  | Bleeding | Yes | 8 | 8 | [03, 14] |
|  |  | No | 89 | 92 | [86, 97] |
|  | Body Malaise | Yes | 62 | 63 | [52, 74] |
|  |  | No | 35 | 37 | [26, 47] |

Even though all participants had gone to the hospital and been diagnosed with Dengue fever they remained poorly informed about the route of Dengue transmission, with 22% of respondents reporting that Dengue fever is spread from one person to another, 30% did not think Dengue is spread by mosquitoes, 2% think that Dengue fever is spread by drinking water and 17% did not know how Dengue fever is spread.

### Prevention from Dengue vector bites

The questionnaire asked about personal protection measures in general – it wasn’t directed towards malaria. Nonetheless, the study found that most respondents think that methods used to prevent malaria vectors work similarly on preventing Dengue vector bites. About 86% of Dengue patients thought that using bed net could prevent Dengue vectors. Other methods mentioned for prevention of mosquito bites were aerosol spray (46%), house screening (27%), Mosquito coils (6%), wearing long dress and trousers (16%) and use of topical repellent 12%, while 20% of participants mentioned the use of fans to prevent mosquito bites, which is ineffective. “Table 4”

### Travel History

A total of 72 (70%) Dengue patients had not travelled out of Dar es Salaam suggesting autochthonous transmission. Of those who travelled, the majority went to other parts of Tanzania, Africa and Asia; 24%, 2%, and 2%, respectively. Two of the Dengue patients recorded at Mwananyamala hospital were not Dar es Salaam residents but they came to Dar es Salaam during the Dengue outbreak. They got sick while in Dar and were treated in Kinondoni District.

## Discussion

Understanding the pattern of Dengue fever and risk factors associated with transmission in Dar es Salaam it is of great significance to prevent future outbreaks through targeted control of Dengue vectors and disseminating appropriate behaviour change communication messages to 1) improve household level environmental modification or destruction of breeding sites around homes, 2) to encourage adherence to personal protection and 3) identification of disease symptoms of the disease as is successfully used in many endemic countries [24]. Future outbreaks have a high probability of occurring more frequently since there have been outbreaks in 2007-8 [7], 2010 (Type III detected in Zanzibar) [25], 2013 (Type II) [26] and 2014 (Type II) [27] as well as locally reported cases in 2017) However, knowledge among communities and about non-malaria febrile illnesses is extremely low and when attending health services, clinical diagnosis for febrile illnesses lacks specificity and has been reported to contribute to misdiagnosis and mistreatment of febrile patients [28].

The study showed that those with higher SES were more likely to have dengue and also more likely to have suitable breeding habitats for dengue vectors around their homes. Those with confirmed Dengue used mainly bednets to prevent mosquito bites, a strategy that does not protect against dengue. Furthermore, knowledge of dengue in this population that had already been diagnosed with dengue was exceedingly low and only 30% of respondents correctly identified dengue transmission as vector-borne. It is generally recognised that people with low SES are usually prone to vector-borne diseases [29-32] including Dengue, although not in all cases as shown by a recent systematic review of the link between Dengue and Socioeconomic status [33]. Our study described the group that is more at risk than others and the spatial distribution of Dengue virus cases has been produced. Because Aedes mosquitoes are known to be prolific feeders with a short flight range they tend to live and feed in a radius of around 100m from a home [14, 34]. In this case, they may bite people in Higher SES households in areas where they are more abundant. Controlling breeding habitats in and around houses of higher-SES is a logical first step for reducing Dengue vector population and the potential spread of Dengue virus. The study results give us a hint that Dengue has not been endemic to Dar es Salaam but as it has a high epidemic potential [35] and therefore environmental management should be the first step in proactive future epidemic prevention.

The study also assessed individual awareness of Dengue infection prevention. The interesting feature of this study was also to create a surveillance tool “habitat suitability-score” that can be used in future years to assess possibilities of Dengue outbreaks before rain season depending on the biology and favourable environment of Dengue vector. The tool demonstrated that higher numbers of cases among high SES groups may be linked to their houses being more suitable for Dengue vector mosquitoes.

Clearly, this tool is only reported here and needs more testing in Dar es Salaam and other sites with *Ae. aegypti* to measure its efficiency for predicting suitable habitats and areas where targeted environmental management should be conducted. However, data agree well with work of Mboera and colleagues [27] in Dar es Salaam and Lutomiah *et al* [36] and Ngugi [37] in Mombasa, who identified large items of household waste such as buckets and tyres to be the most important sources of mosquitoes as was seen here in Kinondoni.

The rainy season amplifies the number of Aedes mosquitoes and unusually heavy rains during climate anomalies such as El Nino are often associated with outbreaks of Dengue [38]. The increase in available breeding sites and consequently of the vectors is likely to have amplified Dengue infection which was likely to be subclinical but to a large number of people large enough to turn into an outbreak [39]. Nonetheless, Dengue cases declined as the rainy season came to an end in July. This may also suggest that outdoor breeding habitats “Fig 6” are more responsible for increasing of Dengue vectors and the spread of Dengue virus into breeding sites than indoor breeding habitats as has also been seen in neighbouring Kenya [37].

**Fig 6.**
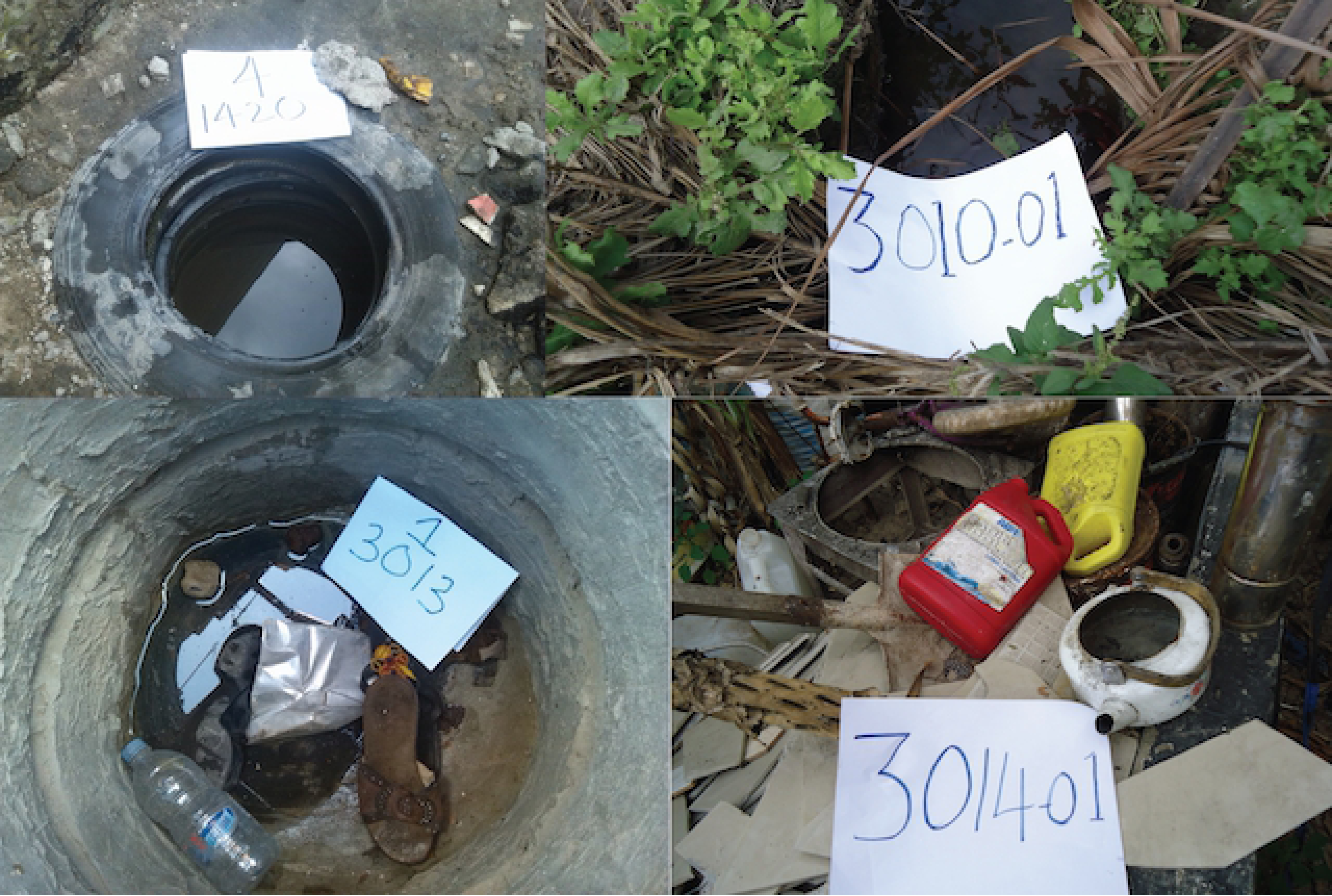
Some of the Breeding Habitat types found around households that had positive Dengue cases traced from Mwananyamala hospital during the 2014 Dengue outbreak.

None of the Aedes found was *Ae. africanus*, which maintains sylvatic yellow fever, or *Ae. albopictus* that is a dengue vector currently invading into a number of African countries [40] but has not yet been found in Tanzania [27, 41]. Study findings show that higher quality breeding habitats in relatively more wealthy households are likely to be a potential source of Dengue vectors. The rainy season may act as an amplifier to the small population of Aedes to reach a threshold adequate to cause an outbreak. Because the accumulation of containers that can sustain Aedes breeding mosquitoes are abundant in market areas due to poor collection of garbage and other wastes, Aedes breeding thrives in and around the market area and people who live and work around these areas may be at increased risk. Since marketplaces in Dar es Salaam are not located in wealthy areas these findings may suggest that the affluent households in poor neighbourhoods could act as a Dengue vector reservoir. Therefore education of more wealthy households in controlling breeding habitats before the rain begins in these houses that reserve water and provide breeding habitat for Aedes mosquito should be prioritized.

### Dengue Public awareness

Most of the people interviewed 60% knew about Dengue when they were already sick. They did not have any information before they got sick. Although Dengue is a neglected disease the outbreak lasted for over six months. People could have received much more information one month to six months earlier during the outbreak but most did not.

On the correlated variable summarized by principal component analysis suggests that using radio to convey disease awareness may have not been a successful means to inform the population simply because there was lower radio information about Dengue fever as it is also reflected in the method from which Dengue patients became aware of Dengue disease “Table 4”. Households’ access to radio was found to be 77% however information about dengue was reported by 38% of the patients. This is almost half of those who have radios. This could be that there were infrequent radio broadcasts and also the time of broadcasting could have missed those at work. Furthermore, television broadcast is only likely to reach the richer population. It is therefore suggested that messages to raise awareness for future outbreaks should be specifically tailored to specific groups by specific medium [42]. For instance, households with dengue patients had mobile ownership at nearly 100% “Table 3”. PCA shows that mobile phones are equally owned across all SES. Therefore, they provide a better platform to disseminate outbreak information to the public. In recent years mobile banking and other financial services through mobile phones have led to the exponential increase of mobile phone ownership which is at 73% in Tanzania [43]. A public health tailored message about an outbreak may efficiently be disseminated through mobile phones.

### Study limitations

This was an observational study in which we were not in control of most variables and we had to rely on information already obtained of dengue patient at the Dengue clinic. This was designated hospital for all Dengue patients in the whole of Kinondoni District. The study took place towards the end of both the Dengue outbreak and the rainy season. Some social-environmental variables had changed. We also cannot establish temporal relationships apart from relying on two weeks of dengue fever incubation period in which we assumed that patients came to the clinic in just a day or two after the onset of the disease. There could also be recall bias from some respondents who were sick several months before we collected information. The study may have missed some Dengue cases that had insufficient discomfort to require medical attention. Another limitation, the study cannot tell if the higher number of people with higher SES among dengue patients is influenced by health-seeking behaviour and affording treatment and have time to attend the clinic.

There was low reporting of Dengue information from radio or television, the study did not ask about the time of broadcasting Dengue information from neither broadcaster nor patients. The broadcasting time could results in a lower number of listeners if it was during work hours.

## Conclusion

Because Dengue outbreaks in Tanzania are related to the rainy season “Fig 3” and market areas were the foci, it is logical for the diseases to outbreak management team to clear the market areas of refuse before the rains start. The intention to prevent Dengue fever should focus on, health care providers at primary care levels, specifically at dispensaries, health centres and municipal hospitals in to detect initial cases of Dengue before they turn into an outbreak. All patients who present with classical signs and symptoms need to be provided with information on how to clear breeding sites from their compounds.

Mass education on vector breeding habitats that involve the management of all containers at the compound that may hold water, frequent emptying and cleaning water-storage vessels, cleaning of gutters; covering stored tyres from rainfall, recycling or discarding of non-essential containers should become a focus for the environmental dengue prevention. It is currently not part of the government vector control strategy which is may benefit from a shift from reactive to proactive vector control [47]. On public awareness and preventive measure, the study advocates the use of mobile phone to disseminate accurate and timely health information. The flourishing of mobile phones ownership and use across all SESs should be a tool to deliver immediate information about dengue preparedness and prevention through information hotlines and tailored text messages.

## Acknowledgement

We thank the commissioners and medical officers of Kinondoni Districts for their cooperation; the medical officers in charge of the Dengue diagnostic centre at Mwananyamala hospital. We thank the data collection team of surveyors’ 1-8 for collecting information and samples from Dengue patient houses. We also thank Dr. Lena Lorenz, Hans Overgaard, William Kisinza and the ABCDR study team as a whole for the comparator data. Lastly but importantly we thank Dr. Naomie Boiloat and her fever project that noticed fever outbreak which happened to be Dengue fever.

## Authors’ contributions

DM, SJM and PC designed the study. DM performed data analysis and wrote the manuscript. TG trained field surveyors and collected vector data. HN and SJM conducted larvae sample and identification at IHI laboratory. YPM analysed GIS data and produced study maps, PC trained field workers and provided entomological tools. SJM sought for study fund supervised data collection and analysis, edited the manuscript.

S1 Checklist: STROBE Checklist

